# Multi-scale spatial heterogeneity enhances particle clearance in airway ciliary arrays

**DOI:** 10.1101/665125

**Authors:** Guillermina R. Ramirez-San Juan, Arnold J. T. M. Mathijssen, Mu He, Lily Jan, Wallace Marshall, Manu Prakash

## Abstract

Mucus clearance constitutes the primary defence of the respiratory system against viruses, bacteria and environmental insults [1]. This transport across the entire airway emerges from the integrated activity of thousands of multiciliated cells, each containing hundreds of cilia, which together must coordinate their spatial arrangement, alignment and motility [2, 3]. The mechanisms of fluid transport have been studied extensively at the level of an individual cilium [4, 5], collectively moving metachronal waves [6–10], and more generally the hydrodynamics of active matter [11, 12]. However, the connection between local cilia architecture and the topology of the flows they generate remains largely unexplored. Here, we image the mouse airway from the sub-cellular (nm) to the organ scales (mm), characterising quantitatively its ciliary arrangement and the generated flows. Locally we measure heterogeneity in both cilia organisation and flow structure, but across the trachea fluid transport is coherent. To examine this result, a hydrodynamic model was developed for a systematic exploration of different tissue architectures. Surprisingly, we find that disorder enhances particle clearance, whether it originates from fluctuations, heterogeneity in multiciliated cell arrangement or ciliary misalignment. This resembles elements of ‘stochastic resonance’ [13–15] in a self-assembled biological system. Taken together, our results shed light on how the microstructure of an active carpet [16, 17] determines its emergent dynamics. Furthermore, this work is also directly applicable to human airway pathologies [1], which are the third leading cause of deaths worldwide [18].

## PATTERNING AND ALIGNMENT OF AIRWAY CILIA ACROSS THE SCALES

The many functions of the airway are reflected by the variety of specialised cell types it contains, including multiciliated, secretory, nerve, immune and basal cells [19]. This inevitably leads to constraints in the organisation of the airway cilia, which is compounded over several orders of magnitude [20, 21] (Fig. 1A). Previous studies have noted the heterogeneity in airway cilia arrangement [22, 23], but they lack quantitative descriptions of disorder across all relevant spatial scales (nm to mm).

**FIG. 1.**
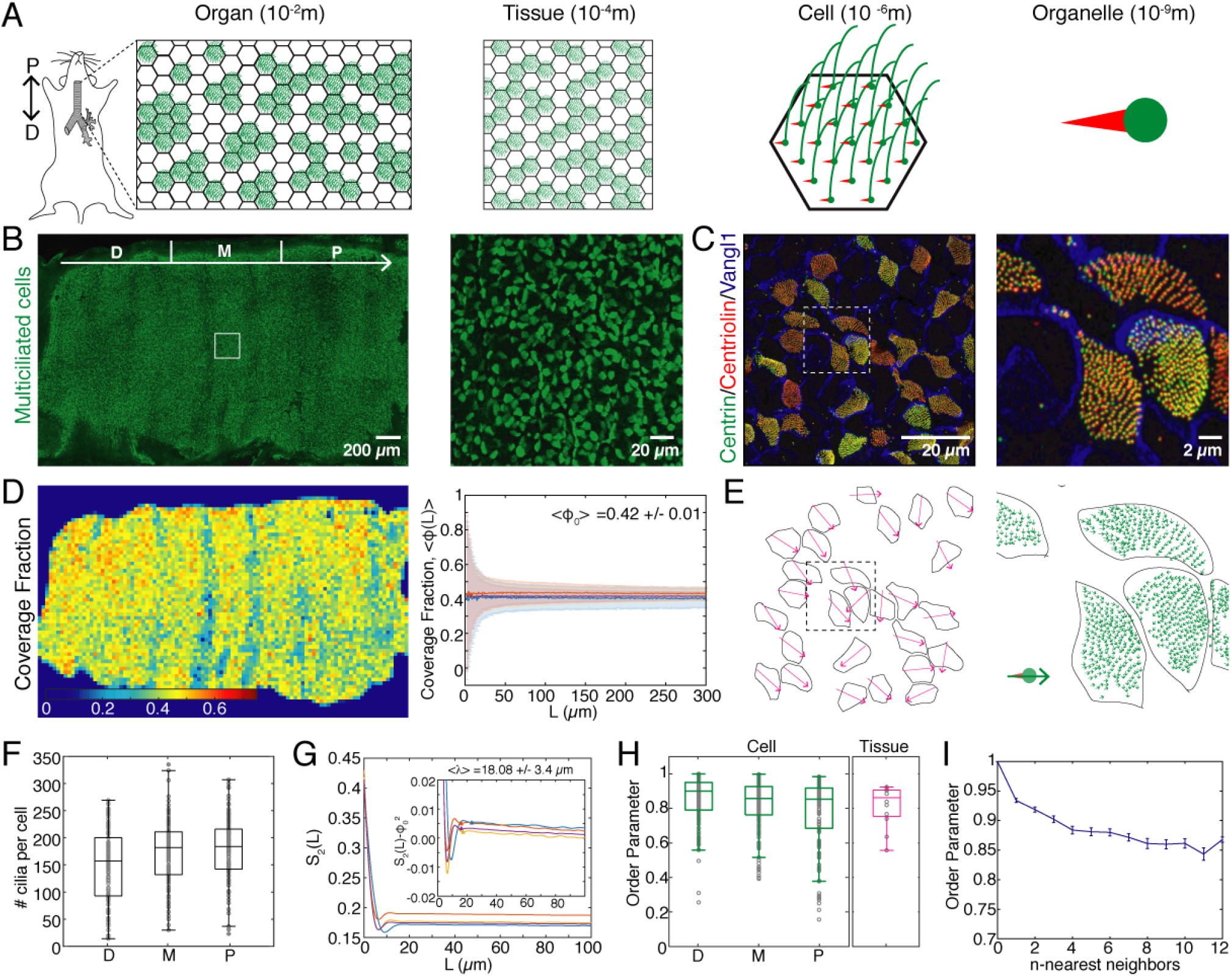
Heterogeneity in spatial patterning of cilia in the mouse airway epithelium. **A**. Schematic of the organisation of multiciliated cells across spatial scales. Thousands of multiciliated cells line the trachea, forming a “patchwork” pattern at the tissue scale. Each multiciliated cell contains hundreds of cilia anchored to the cell by the basal body. **B**. Fluorescence image of an entire trachea where multiciliated cells are labelled by Centrin-GFP. The distal, medial and proximal (D|M|P) segments of the organ, each one third, are indicated by white separation marks. The direction of fluid transport is D→P (lung→oral). Right: Magnification of region outlined by the white square. **C**. Fluorescence image of a section of the multiciliated epithelium. Vangl1 localizes to cell membranes. Fluorescently labelled Centrin and Centriolin mark individual basal bodies. The vector that connects Centriolin→Centrin defines the orientation of an individual cilium. Right: Magnification of region outlined by the white square. **D**. Heat map of the coverage fraction of multiciliated cells (*ϕ*). Right: Coverage fraction measured in randomly selected square windows of size *L*, uniformly distributed along the trachea. 〈*ϕ*_0_〉 is the average over all *L* and all tracheas imaged. **E**. Tissue scale orientation field obtained by averaging the orientation of cilia within each multiciliated cell shown in C. Right: Orientation of individual cilia shown in panel C. **F**. Quantification of the number of cilia per multiciliated cell in the distal (D), medial (M) and proximal (P) segments of the trachea. **G**. Plot of the two-point correlation function. Inset: Wavelength of the patchwork pattern formed by multiciliated cells. Triangles mark the point at which oscillations in 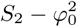 vanish. **H**. Orientational order parameter m calculated for multiciliated cells in the D,M,P segments of the trachea (green) and tissue-scale orientational order parameter M (magenta). **I**. Average orientational order parameter as a function of the size of the tissue region analysed. The size of each region grows from encompassing adjacent neighbours to n-connected regions.

To quantitatively map the airway architecture across these scales, we imaged transgenic mice expressing a fluorescently tagged version of Centrin (multiciliated cell, basal body specific marker). Along the distal-proximal (*D*-*P*, lung-oral) axis of the trachea, multiciliated cells cover only a fraction of the surface, corresponding to *ϕ*_0_ = 0.42 ± 0.01, giving rise to a ‘patchwork’ pattern (Fig. 1B; Fig. S1A). Multiciliated cells are distributed uniformly along the *D*-*P* axis, except at the cartilage rings where their coverage fraction decreases (Fig. 1D; Fig. S1A).

To characterise the structure of the multiciliated cell pattern, we measured its periodicity by computing its two-point correlation function *S*_2_(*L*) (see Methods § 2 a). From this function we obtained the correlation length, the wavelength *λ* = (18.08 ± 3.4)*μ*m, which corresponds to approximately 4 cells, defined by the distance at which the oscillations in the function 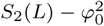 disappear (Figure 1G).

Furthermore, measurements from fixed tissues agree with the values obtained for the coverage fraction *ϕ* = 0.41 ± 0.04 and wavelength *λ* = (25.52 ± 7.35)*μ*m measured from live trachea imaging (Fig. S1B; Suppl. Video 1). The wavelength and coverage fraction can be measured with higher accuracy on fixed tissues. However, we analysed time-lapse videos of the tissue to confirm that all multiciliated cells are active in the trachea. Indeed, agreement in measurements from fixed and live data shows that all multiciliated cells contribute to the flow generated by the carpet (Fig. S1C,D).

Each multiciliated cell is decorated with hundreds of cilia [21, 24]. Quantification of the number of basal bodies from images of fixed trachea segments (Fig. 1C) showed that each multiciliated cell has on average *N*_*cilia*_ = 169 ± 61 cilia. We note that this number fluctuates strongly between 20-350 cilia per cell (Fig. 1F).

Individual cilia beat with an asymmetric wave form confined to a plane [25], with a polarity vector that is established during organismal development through cytoskeletal arrangements and hydrodynamic interactions, that becomes fixed at adulthood [20, 21, 26]. This beating plane is defined uniquely by the polarity of the basal body [21], which is marked by Centrin, and its basal foot, which we marked with Centriolin. Hence, for each cilium *i* we establish its orientation vector ***p***_*i*_ by connecting the relative positions of its Centrin and Centriolin puncta (Fig. 1E; right panel, green arrows). Subsequently, for each multiciliated cell *j* we average its cilia vectors to define the average orientation of cilia within a given cell ***P***_*j*_ (Fig. 1E; left panel, magenta arrows).

From these vectors we observe that cilia exhibit significant fluctuations in their relative orientation, both across the tissue and within individual cells. These fluctuations are quantified by the order parameter, 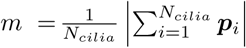, analogous to magnetisation. We find an average order parameter of 〈*m*〉 = 0.81 ± 0.08, representing the orientational heterogeneity at the subcellular scale. At the tissue scale the order parameter is 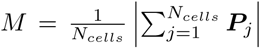, where *N*_*cells*_ is the number of cells. Here, we find 〈*M*〉 = 0.81 ± 0.12, with small variations across different tissue sections (Fig. 1H). Interestingly, the average order parameter decays with distance, measured in terms of number of neighbouring cells considered (Fig. 1I), indicating that cells are more aligned with their immediate neighbours compared to cells that are further away in the tissue. Altogether, these results show that heterogeneity is a feature of the spatial organisation of airway cilia from the subcellular to the organ scale.

## CILIARY FLOWS ARE GLOBALLY COHERENT BUT LOCALLY HETEROGENEOUS

Perfectly aligned and densely covered ciliary arrays will generate directed flows [27], but our results show that the trachea inherently features patchiness and misalignment (Fig. 1). In particular, this patchy architecture raises the key question of how particle transport is connected over regions devoid of cilia. Thus, we next investigated the flows generated by multiciliated cells in the trachea.

To monitor flow structure, fluorescent beads were added to the media as tracer particles. We find that flows across entire tracheas are coherent and globally directed along the *D*-*P* axis (Fig. 2A,C left panel; Fig. S2 A; Suppl. Video 2) with an average speed of (8.4 ± 2.6)*μ*m/s (Fig. 2E). However, at a scale comparable to the wavelength of the multiciliated cell “patchwork” pattern, *λ* = (18.08 ± 3.4)*μ*m, we observe variations in the local flow direction and magnitude (Figure 2A,C right panel; Fig. S2B; Suppl. Video 3). Specifically, we see connected regions with stronger flow, separated by regions of weaker flows. To quantify the structure of these currents we calculate the spatial, *δC*_*vv*_(*R*), and temporal *δC*_*vv*_(*τ*) correlations (see Methods § 2 d). For each of our measurements, *δC*_*dd*_(*R*) decays exponentially over 30*μm* and can be fitted well in this range by an exponential, *δC*_*dd*_(*R*) = *e*^(*−R/R*0)^ (Sup. Fig. S2C), which defines the correlation length *R*_0_. Thus, the domain size of correlated flow is *R*_0_ = (13.7 ± 5.5)*μ*m (Fig. 2H), which is comparable to the wavelength *λ* of the pattern of multiciliated cells. Moreover, the timescale of temporal decorrelation, *τ* = (0.029 ± 0.01)s (see Methods § 2 d) is on the same order of magnitude as the timescale of the cilia beat frequency, *CBF* = (11.8 ± 2.3)Hz (Fig. S2D, E; Suppl. Video 4). Therefore, the spatiotemporal flow structure coincides with the spatiotemporal organisation of the heterogeneous ciliary carpet.

**FIG. 2.**
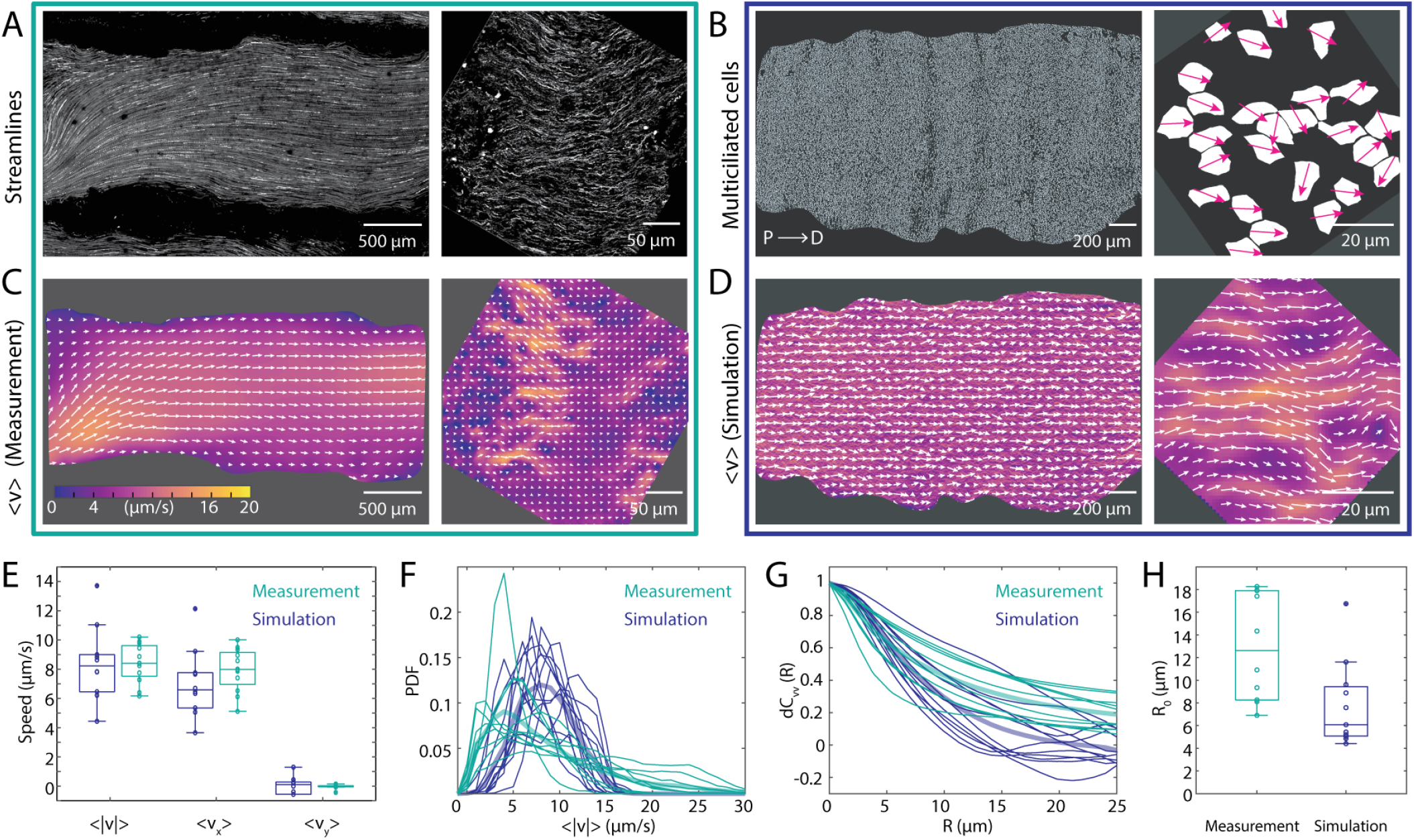
Measurements of ciliary flows at the tissue and cellular scales. **A.** Pathlines of the flow generated by multiciliated cells across the entire trachea (left). Right: Pathlines of the flow visualized at a scale comparable to the wavelength of the cilia patchwork pattern. **B.** Binary images obtained from thresholding the images of multiciliated cell localization shown in Fig. 1B,C (left panels). White regions represent multiciliated cells. Right: Magenta arrows show the average orientation of cilia within each multiciliated cell measured from Fig. 1C. These coverage and orientation fields are used as input for the model, shown in panel D. **C.** Representative experimental flow fields measured at the scale of the entire trachea (left) and with micrometer resolution (right). Colours show the flow velocity magnitude and white arrows indicate the streamlines. **D.** Simulated flow fields, using the experimental data of panel B as an input.Colours show the flow velocity magnitude and white arrows indicate the streamlines. **E.** Comparison of flow speeds between experiment (green) and model (blue). The majority comes from the x component. Boxplot central mark indicates the median, and the bottom and top edges of the box indicate the 25^*th*^ and 75^*th*^ percentiles, respectively. **F.** Comparison of the flow magnitude distribution, PDF(〈|*v*|〉), between experiment and model. Thin lines show the PDF from a field of view of the size shown in C,D (right). Thick lines are the average of all flow fields analyzed. **G.** Comparison of the spatial autocorrelation function between experiment and model. Thin lines show measurements from a field of view of the size shown in C,D (right). **H.** Comparison of correlation length *R*_0_ from the experiment, 〈*R*_*0*_〉 = (12.94 ± 4.7)*μ*m, and from the model, 〈*R*_*0*_〉 = (7.77 ± 3.8)*μ*m.

To understand how globally directed flows emerge from micron-scale fluctuations, we develop a hydrodynamic model inspired by the ‘envelope approach’ introduced by Lighthill and Blake (see Methods § 3). Multiciliated cells are represented by a slip velocity *U* at the tissue surface that follows the orientation of the underlying cilia. The 3D flow is then computed using a computational fluid dynamics (CFD) solver.

First, we simulate the flows using as input the global and local tissue configurations measured from mice tracheas (Fig. 2B). In agreement with our experiments, we observe globally coherent currents directed across the trachea, but at the micrometer scale we see heterogeneity in the magnitude and direction of the flows (Fig. 2D). Comparison of the statistical properties of simulated and measured flows, shows that flow velocity distributions and their correlation lengths agree, establishing the validity of our model (Fig. 2E-H).

## FLOW STRUCTURE IS DETERMINED BY MULTICILIATED CELL ARCHITECTURE

Our measurements of the flow generated by multiciliated cells in the trachea show that complete orientational order and coverage are not a requirement for a ciliary carpet to generate directed fluid flows. However, it is unclear how much variability in multiciliated cell configuration can be tolerated before fluid transport is impaired. Therefore, we next use our model to uncover how changes in the spatial distribution of multiciliated cells impact the properties of the flow they generate (Fig. 3A,B).

**FIG. 3.**
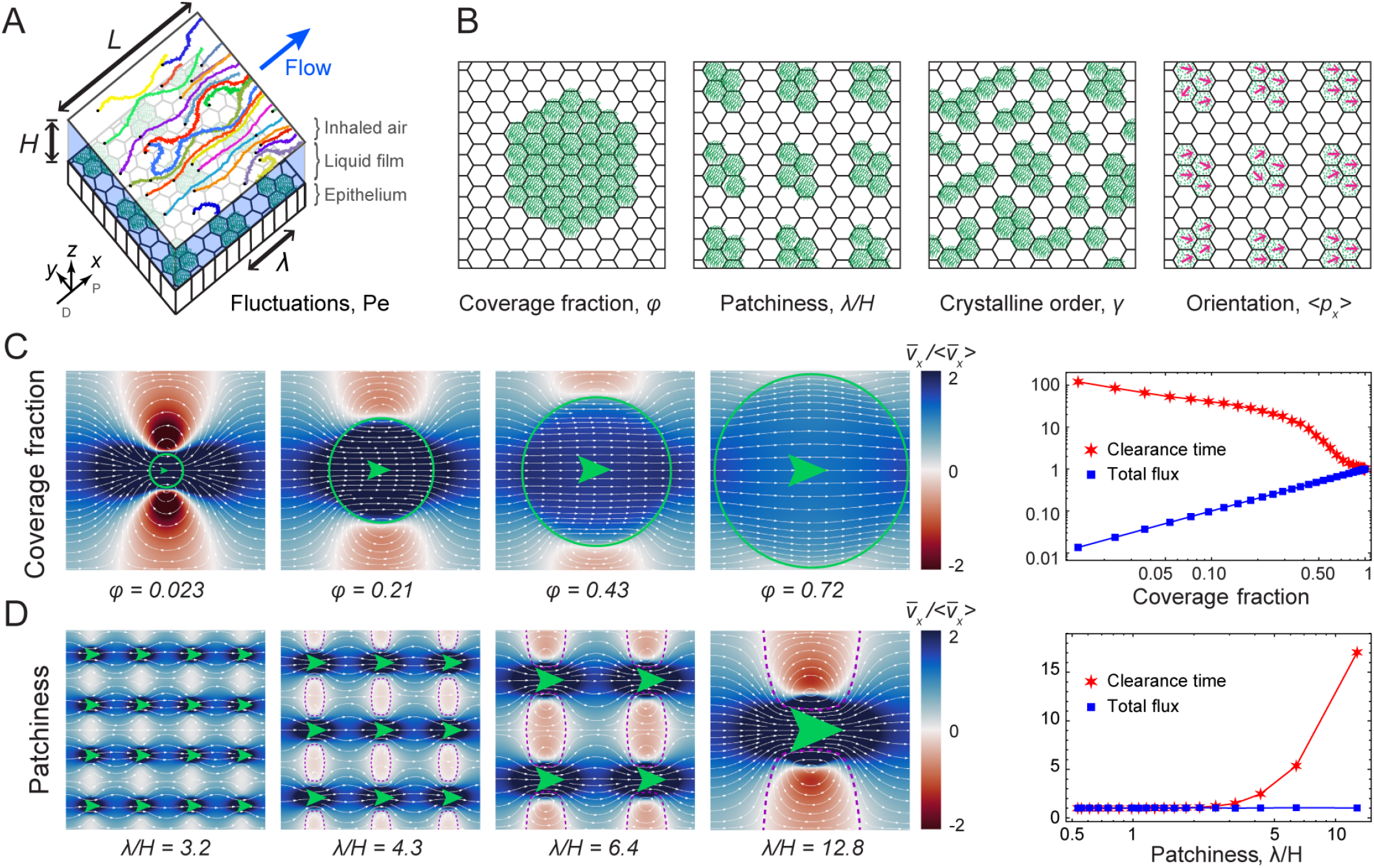
Hydrodynamic model for particle flux and clearance. **A.** Diagram of simulation geometry. A pattern of multiciliated cells (green) drives a liquid film flow (blue) in a 3D computational fluid dynamics (CFD) simulation with periodic boundary conditions in *x* and *y*. Particle trajectories with random initial positions are subject to this flow and fluctuations. **B**. Total flux and clearance times are examined as a function of ciliary organisation, characerised by the coverage fraction *ϕ*, wavelength *λ*, crystallinity *γ*, and orientation order 〈*p*_*x*_〉. **C**. Higher coverage fractions improve clearance. Left: Shown are streamlines (white) and the longitudinal flow strength (red-blue) as a function of coverage fraction *ϕ*. Green circles indicate the area covered by cilia and green arrows their orientation. Right: Corresponding plot of clearance time and total flux, normalised with respect to full coverage, showing the mean of an ensemble of *N*_*p*_ = 10^4^ particle trajectories. **D**. Patchiness induces recirculation and slows clearance. Left: Flows as a function of wavelength *λ*. Dashed purple lines are separatrix streamlines that isolate recirculation zones (red) from the main currents (blue). Right: Corresponding clearance time and flux, normalised with respect to uniform coverage, again averaged over *N*_*p*_ = 10^4^ particle trajectories.

We explore systematically how total flux and particle clearance time change as a function of variations in multiciliated cell coverage fraction, wavelength, and orientational and geometrical order. The total flux *J* = ∫ *v*_*x*_*dydz* measures the strength of the flow ***v***(***r***) globally. To account for the local flow microstructure, we compute the mean first passage time 〈*T*〉 taken to travel a distance *L* along *x* for an ensemble of particles advected by the flow (See Methods § 3 d). This is a measure of streamline connectivity, thus flow topology, rather than only flow strength.

First we test the role of coverage fraction on flow architecture. We observe that the total flux generated by the ciliary carpet increases linearly with coverage fraction (Fig. 3C, right panel). In the limit of low coverage (*ϕ* → 0) the flow structure features recirculating eddies, approaching the flow due to a point force confined in a liquid film or Stokeslet (see Methods § 3 a). As the coverage fraction grows, regions of negative flux disappear and in the limit *ϕ* → 1 uniform flow is recovered. Consequently, the clearance time (red stars) increases at low *ϕ*, as expected, but not linearly. A reduction of coverage therefore strongly impacts flow strength.

We then vary the periodicity of the multiciliated cell pattern. We define “patchiness” as the dimensionless number *λ/H*, where *λ* is the wavelength of the multiciliated cell pattern and *H* is the height of the fluid film. We keep the coverage fraction and film height constant, distributing the same amount of multiciliated cells in clusters of different sizes (green arrows), varying only the wavelength *λ* (Fig. 3D). The total flux *J* is constant for all configurations, because the total force exerted on the liquid is the same. However, the streamline structure changes as a function of multiciliated cell arrangement. For large patchiness recirculating eddies emerge, creating zones where particles are trapped in closed streamline orbits (dashed purple separatrices). However, when cells are distributed more homogeneously, these zones disappear and consequently clearance times are reduced (Fig. 3D). We note that the emergence of recirculation zones can be tuned independently by reducing the film height. Therefore, higher patchiness caused either by dehydration of the liquid film or long wavelengths leads to impaired clearance.

## DISORDER CAN ENHANCE PARTICLE TRANSPORT

Having examined how deterministic parameters of multiciliated cell organisation influence flow structure, we now analyse the impact of introducing disorder in these ciliary arrays. Surprisingly, we find that different types of disorder can improve clearance.

In the trachea, patches of multiciliated cells lack crystalline order (Fig. 1B,D). We introduce this geometric heterogeneity by shifting the positions of ciliated patches along *x* and *y* with standard deviation *σ*. Thus, geometric heterogeneity is described by the order parameter 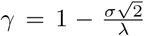, the crystallinity. The resulting flows (Fig. 4A) again feature a constant total flux but, importantly, the geometric disorder breaks open the closed orbits, causing streamlines to cut through the recirculation zones (black dashed streamlines), and strongly reducing clearance time 〈*T*〉.

**FIG. 4.**
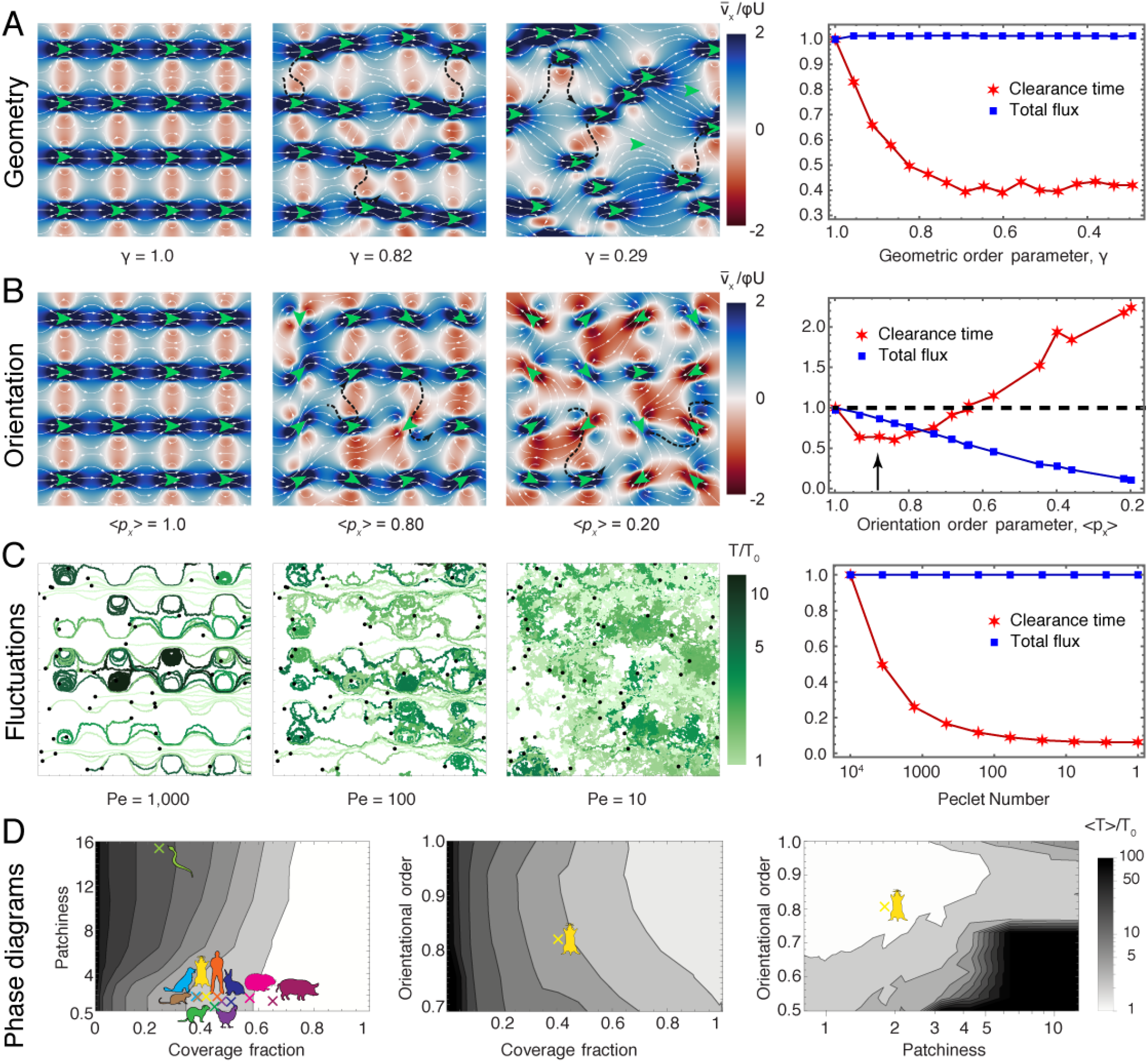
Disorder improves clearance in heterogeneous epithelia. **A**. Left: Flow strength along the *D-P* axis (red-blue) and streamlines (white) as a function of crystallinity *γ*. Green arrows indicate the position and orientation of the cilia patches. A few typical (black dashed) streamlines are shown to cross through recirculation areas. Right: Corresponding total flux and clearance time, normalised with respect to crystalline order. *N*_*p*_ = 10^3^ particle trajectories. **B**. Flow and clearance as a function of cilia orientational order 〈*p*_*x*_〉. Same as (A) otherwise. **C**. Left: Particle trajectories subject to flow and diffusion for a range of Péclet numbers, Pé = *λϕU/D*. Right: Corresponding total flux and clearance time, normalised with respect to weak fluctuations. *N*_*p*_ = 10^4^ particle trajectories. **D**. Phase diagrams showing clearance time (grey scale, defined in legend) for the combinations of (*ϕ, λ/H, M*), with constant *M* = 1, *λ/H* = 8 and *ϕ* = 0.4, respectively. The mouse cartoon indicates the measurements for the mouse epithelium. Other coloured cartoons on the first panel show values of patchiness and coverage fraction measured for several different animals, from data available in the literature (See Methods § 2 f and Figure S4).

Next, we examine disorder in the cilia orientations (Fig. 1H), by varying the orientational order parameter 〈*p*_*x*_〉 = *M* (Fig. 4B). The total flux is no longer conserved and decreases with orientational noise. Interestingly, we observe a biphasic behaviour in clearance time: for weakly misaligned cilia the recirculation zones are broken up and the clearance time is reduced, yet, the clearance time increases for strong misalignment. We predict an optimum around *M* = 0.89 ± 0.06 (black arrow). Looking back at measured values, *M* = 0.81 ± 0.12 (Fig. 1H), we see that the mouse tracheas occupy the same region in this parameter space.

Finally, we study the effect of diffusion, *D*, which can be due to thermal or ciliary beating noise (Suppl. Video 5). To understand how diffusion impacts particle clearance, we compare the advection time *τ*_*a*_ = *λ*/(*ϕU*) with the diffusion time *τ*_*e*_ = *ε*^2^/*D* required for particles to escape eddies of size *ε* ∼ *λ*. The ratio of these times defines the Péclet number, Pé = *τ*_*e*_/*τ*_*a*_ and their sum gives the mean clearance time, 〈*T*〉 ~ *τ*_*e*_ + *τ*_*a*_. Therefore, with weak fluctuations (Pé ≫ 1) the particles remain stuck in closed orbits (Fig. 4C, left), but strong diffusion (*τ*_*e*_ → 0) reduces clearance times by helping particles escape, and connecting regions of closed streamlines (Fig. 4C, right). We note that diffusion only reduces clearance time in carpets with closed eddies, e.g. at high patchiness, since *τ*_*e*_ = 0 for all values of Pé in the absence of recirculation zones.

To cast our results in a broader context, we map the phase space of clearance times in terms of ciliary organisation (Fig. 4D). First, we observe that for a large range of coverage fractions (*ϕ* ≳ 0.58) the clearance time is minimal. Surprisingly, the mouse airway operates below this ideal regime, but its low patchiness and moderate disorder in cilia orientation still allow for clearance times to be relatively small. This observation raises the question whether airway tissues from other animals have similar, sub-optimal, coverage fractions. A brief literature survey showed that different species have similar coverage fractions and patchiness (Figure S4), clustering in the same region of phase diagram as the mouse (Figure 4). Snakes form an exception, potentially because of their unique trachea morphology. Given that patchiness is conserved among organisms with drastically different trachea length, we conjecture that maintaining this number fixed among different species provides an effective mechanism for clearance. However, further studies comparing the morphology of tracheas will be required to understand whether this design principle is universal.

## DISCUSSION

Our work establishes a quantitative link between the topology of the flow generated by a ciliary array and its underlying spatial organisation. We investigate the arrangement of cilia and their associated flow generation, from the molecular to the organ scale, in the mouse conductive airway. Using a hydrodynamic model we explore the effect on flow topology on varying ciliary arrangements. Making the important distinction that particle clearance is not necessarily proportional to the overall liquid flux, but rather set by streamline connectivity, we discover that disorder facilitates globally connected particle clearance. This disorder arises from heterogeneity in ciliary arrangement, ranging from the patterning and alignment of individual cilia, up to the arrangement of multiciliated cells in the entire trachea.

Even if surprising, it is not uncommon in physical and biological systems that ‘noise’ can prove beneficial, a concept sometimes referred to as stochastic resonance (SR) [13–15]. In these systems noise levels are initially advantageous but often only up to a point, the resonance peak, after which more fluctuations become unfavourable [13]. The organisation of cilia should be understood in this context of trade-offs in self-assembled systems. Indeed, across diverse physiological contexts, disorder in the spatio-temporal organisation of cilia can be harnessed to optimise their biological function. For example, dynamic switches in beating patterns modulate the transport of cerebrospinal fluid [28], length differences can create hydrodynamic microhabitats [29], and topological defects can enhance feeding in starfish larvae [16] and attract nutrients to active carpets [17].

However, disorder can also be detrimental to physiology, as reduced ciliary coverage and alignment are a hallmark of airway pathologies [30, 31]. Our phase diagrams (Fig. 4D) provide a framework to understand how much heterogeneity can be tolerated before clearance times increase beyond a critical value. Understanding the link between ciliary organisation and flow structure could open up new avenues for diagnostics and therapies for airway pathologies.

Beyond understanding airway physiology, our phase diagrams (Fig. 4D) prescribe the design principles for synthetic active carpets. This could open new doors to fabricate active surfaces that drive fluid flows with programmable and adaptive topologies, with diverse engineering applications including microseparation and cellular rheometry.

## MATERIALS AND METHODS

### 1. Tissue imaging

#### a. Mouse husbandry

Arl13b-mCherry; Centrin2-GFP (JAX 027967,[32]) were maintained in FVB/N background. This mouse strain was used for all imaging performed. All mice imaged where at stage*>*p15, to ensure cilia orientation is fixed [22]. Mice were housed in an animal facility and maintained in a temperature-controlled and light-controlled environment with an alternating 12-h light/dark cycle. A maximum of five mice were housed per cage. All protocols have been approved by the University of California San Francisco Institutional Animal Care and Use Committee.

#### b. Tissue staining

Mice were euthanized in accordance to the University of California San Francisco Institutional Animal Care and Use Committee. Tracheas were dissected out after euthanasia. To maintain the integrity of the trachea, dissection cuts were performed right below the cricoid cartilage on the proximal end, and below the carina on the distal end.

Tissues were mounted as follows: For whole mount tissue staining, tracheas were sliced open along the proximal-distal axis. For basal body staining, tracheal segments (C1-12) were dissected and sliced open along the proximal-distal axis.

Dissected tissue was fixed with cold methanol for 10 minutes. After fixation, samples were washed and blocked with IF buffer (1x PBS with 1% heat-inactivated goat/donkey serum and 0.3% Triton X-100). Primary antibodies were added and incubated for 1 h at room temperature or overnight at 4 °C. After washing with IF buffer, secondary antibodies and DAPI were added for 1 h at room temperature. Samples were washed with 1x PBS and mounted with Fluoromount-G (SouthernBiotech).

Antibodies used for immunofluorescence staining are mouse monoclonal anti-Centriolin (Santa Cruz: sc-365521; 1:500 dilution) and rabbit anti-Vangl1 (Sigma: HPA025235; 1:1000 dilution), and Alexa Fluor 594-goat anti-rabbit and 633-donkey anti-mouse conjugated secondary antibodies (Invitrogen; 1:1000 dilution).

#### c. Microscopy and live-tissue imaging

Whole-mount fixed tracheal samples were imaged using a Zeiss LSM 780 laser scanning confocal microscope and a 20X Plan Apo NA 0.8 air objective (Stanford Cell Sciences Imaging Facility). The entire surface of the trachea was broken into tiles for imaging. For each tile a z-stack with height of 20-30*μm* was obtained. Images processed for analysis and shown in figure 1B correspond to the maximum intensity projection of z-imaging for each tile.

To image basal body orientations a Zeiss LSM880 laser scanning confocal equipped with Airyscan and 40X Plan Neo NA 1.3 oil immersion objective was used (Gladstone Institutes Histology and Microscopy Core). For each image a z-stack of 5-10 *μm* in heigh was taken. Images processed for analysis and shown in figure 1C correspond to the maximum intensity projection of z-imaging.

For flow imaging, tracheas were dissected in ice-cold Dulbecco’s Modified Eagle Medium: Nutrient Mixture F-12 (ThermoFisher, DMEM/F12), and sliced open along the proximal-distal axis. Samples were kept on ice prior to live imaging. Tracheas were mounted as follows: Two strips of parafilm were attached to a 35mm cell imaging dish (MatTek) to serve as spacers. A single trachea was placed with its multiciliated surface facing the glass bottom of the dish and covered with a 18mm round coverslip. The media surrounding the trachea consisted of a solution of fluorescent beads (2 or 0.5 *μm* diameter; FluoSpheres Invitrogen, 1:250 to 1:500 dilution) in 500*μ*l DMEM/F12 media. Imaging was performed in an environmental chamber (5% CO_2_ at 37 °C) with an inverted microscope (Ti-E; Nikon) equipped with a confocal scan head (Andor Borealis CSU-W1), laser merge module containing 405-,488-, 561-, and 640-nm laser lines (Andor 4-line laser launch) and Zyla 5.5 s-CMOS camera (Andor, Oxford Instruments). Images of global and local flows were acquired using 4X Plan Apo NA 0.2 dry objective and 40X Plan Apo LWD NA 1.5 water immersion objectives.

For ciliary beat imaging, pieces of tissue containing 2-3 tracheal segments were dissected in ice-cold Dulbecco’s Modified Eagle Medium: Nutrient Mixture F-12 (ThermoFisher, DMEM/F12). Tissue sections were sliced open along the proximal-distal axis from the dorsal side and mounted onto 35mm cell imaging dishes (Mat-Tek), between 2 strips of parafilm that served as spacers and covered with an 18mm coverslip. Tissues were immersed in 500ml DMEM/F12. Imaging was performed in an inverted epifluorescence microscope equiped with an environmentally controlled chamber (5% CO_2_ at 37 °C), light source (Sutter XL lamp), DIC optics and a Hamamatsu Flash4.0 camera. Images were acquired using 40X DIC Plan Apo LWD NA 1.5 water immersion and 60X DIC Plan Apo VC NA 1.4 Oil immersion objectives.

### 2. Image analysis

#### a. Analysis of cilia coverage fraction and wavelength

The coverage fraction and wavelength of the multiciliated cell pattern were calculated from fixed whole mount tracheas. Image tiles corresponding to an entire trachea were stitched using the ImageJ “Stitching” plugin [33]. Subsequently, stitched images were binarized by applying a band pass filter and the “imbinarize” function in MATLAB.

To minimize artifacts of thresholding, we removed objects with area*<*30 pixels (12*μm*^2^, for reference a typical cell has an area of ~25*μ*m^2^). For every binarized image the coverage fraction, *ϕ* was measured for 1000 square tiles of length *L*. Tiles were distributed uniformly along the entire image and were chosen randomly. The coverage fraction for each tile is defined as:

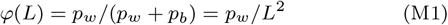

where *p*_*w*_ and *p*_*b*_ are the number of white and black pixels respectively. Thus the average coverage fraction computed from every image is:

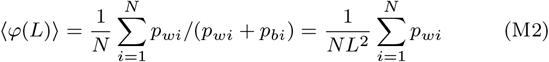

To extract the wavelength of the pattern, we considered the two point correlation function of the binarized image, *f*:

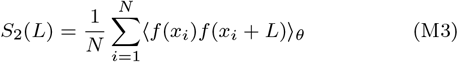

where *x*_*i*_ are all the pixels in the image, N is the total number of pixels in an image and the angle brackets indicate a spatial average along the axes parallel to the sides of a square. We note that *S*_2_(0) = *ϕ*_0_. From this function we obtain the correlation length, or wavelength *λ*, defined by the distance at which the oscillations in the function 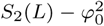 dwindle to zero, or in practice, within 10^*−*3^ of zero [34].

All calculations were performed using custom-written routines in MATLAB.

#### b. Analysis of cilia activity

Additional measurements of coverage fraction and wavelength were performed from time-lapse data of cilia activity as follows.

We identified regions of cilia activity from image stacks that correspond to 1-10 seconds of imaging captured at a frequency *>*100fps. The median image of the timelapse was obtained and substrated from all images, from this resulting stack a maximum intensity projection was obtained and the resulting image was thresholded. This method allowed for identification of regions where pixel values fluctuate due to ciliary beating, rendering a binarized image where the white pixels correspond to regions of cilia activity (See figure S1C). From thresholded images, the two-point correlation function was calculated according to M2 and M3. The coverage fraction, *ϕ*, is *ϕ*_0_ = 〈S_2_(0)〉 and the wavelength, λ, is the first value at which the amplitude of oscillations in 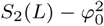 are less than 10^−3^.

Image thresholding was done in ImageJ and calculations of λ and ϕ were performed using custom-written routines in MATLAB.

#### c. Analysis of number of cilia and orientations

The number and orientation of individual cilia was calculated from fixed tissue samples where Centrin and Centriolin were labelled. Individual basal bodies were identified from Centrin images using the “bpass.m” and “pkfnd.m” functions from the Daniel Blair and Eric Dufresne MATLAB version of the particle tracking code in [35]. This yielded the position of the center of each punctate structure, corresponding to a basal body, with sub-pixel accuracy.

To calculate the average number of cilia per cell, the boundaries of cells in an image were traced manually from Vangl1 staining. Basal bodies were binned by cell and counted to obtain the number of cilia per cell *N*_*cilia*_, shown in Figure 1F.

The orientation of each cilium was obtained as follows. First, images were binned in windows of 50×50 pixels centered around identified basal bodies. The built-in “xcorr2 function was applied to calculate the 2D cross correlation of each window in a Centrin image with the same spatial window in the Centriolin image. A unit vector connecting Centrin→Centriolin in the direction of the maximum correlation between images was drawn for every basal body identified.

This yielded a vector field for each image that indicates the orientation of individual cilia. In some cases Centriolin staining was weak, thus the “xcorr2” function yielded a vector in a random direction that did not correspond to the orientation of the basal body. To correct for this, vectors were filtered by calculating the average orientation of cilia in bins of 20 nearest neighbours. Vectors that formed an angle of more than 53° with the mean orientation vector were excluded from the orientation field.

The cellular-scale order parameter, *m*, was computed by binning the filtered orientation vectors by cell, ***p***_*i,j*_ for cilium *i* in cell *j*, and calculating:

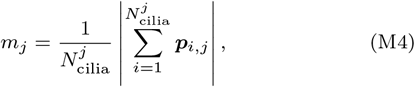

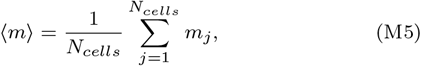

where 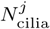 is the number of cilia in cell *j*. Values of 〈*m*〉 for all cells analysed are shown in Figure 1H grouped by proximal, medial or distal, based on the region of the trachea an image was obtained from.

To calculate the tissue-scale order parameter we consider the vectors that correspond to the average orientation of cilia within individual cells:

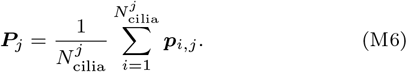

This mean orientation was obtained for all cells in an image. For every image the tissue scale order parameter, *M*, was calculated according to:

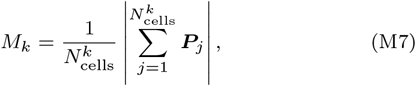

where 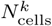 is the number of cells in image *k*, so the average over *N*_*images*_ gives

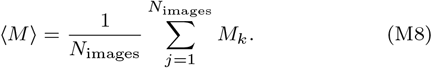

All values obtained are shown in Figure 1H (pink box). The same definition was used to calculate the order parameter for regions of the tissue that include only cells that are n-nearest neighbours (Figure 1I).

All calculations were performed using custom-written routines in MATLAB.

#### d. Analysis of flows

Flow fields were obtained from time-lapse imaging of tissues were fluorescent beads were added to the media. Particle Image Velocimetry (PIV) fields were generated using mpiv (http://www.oceanwave.jp/softwares/mpiv/) for MATLAB. Field parameters were optimized based on bead dilution used in flow field visualization experiments, and then held constant across all analyses to reduce systematic errors due to inconsistent discretization. Parameters used were interrogation window=32 pixels, overlap window=16 pixels, extension window=128 pixels. Temporal parameters for images captured with a 4X objective were temporal averaging= 5 minutes and time step=1 second. For images captured at 40X magnification, parameter values used were temporal averaging= 1 second and time step=0.3-0.13 seconds.

Probability density functions (PDFs) were obtained for fields measured from tissues. To assign units to the simulated data, the average speed for all simulated flows was matched to the average of the speed in ten flow fields measured.

To characterise the temporal and spatial structure of the flow we calculated the spatial and temporal correlations defined as follows:

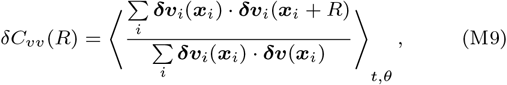

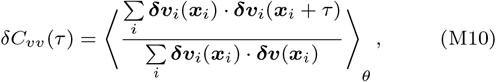

where *δ****v***(*x*) = ***v*** − ***v*** is the flow velocity fluctuation field at position ***x***, and *t*, *θ* indicate temporal and spatial averaging. Since we are interested in understanding flow fluctuations, rather than autocorrelating the velocity fields, ***v***, we first subtract the mean velocity vector of each frame, 〈*v*〉, and autocorrelate the velocity fluctuation field, *δ****v***_*i*_.

We calculate the correlation lengths by fitting an exponential *e*^−*R*/*R*_0_^ or *e*^−*R*/*τ*_0_^ to the correlation functions using the matlab function fit toolbox. The correlation length is defined as *R*_0_ and the temporal correlation time by *τ*_0_.

#### e. Analysis of ciliary beating

The cilia beat frequency and wave velocity were measured from time-lapse imaging of ciliary beating with a temporal resolution of 250-350 frames per second.

Kymographs were obtained for lines drawn parallel or perpendicular to cilia in an image (Figure S2E). Lines were drawn along traces in the kymograph that correspond to cilia beat cycles. The slope of these lines was averaged over all lines in a kymograph, yielding a measurement of cilia beat frequency (parallel to cilium, blue line) or group wave velocity (perpendicular to cilium, green line).

#### f. Literature survey of patchiness and coverage fraction

We conducted a literature survey to find images of the airway tissue where the pattern formed by multiciliated cells could be identified.

Scanning electron microscopy images were found in the literature for airway tissue from the following animals: Chicken [36], dog [37], ferret [38], hamster [39], human [40], pig [41], rat[42] and snake [43].

Images were cropped from figures, to include only the region where the multiciliated tissue was present (Figure S4). Regions of the image containing multiciliated cells were identified and drawn manually. This generated a binary image where the white pixels corresponded to areas where multiciliated cells are present. Coverage fraction and wavelength were calculated from binary images as described in § 2 a. Coverage fraction measurements were reliable, however, given the limited number of pixels in the image, measurements of the two-point correlation were noisy, thus the wavelength was estimated to be the the value at which 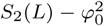 is zero or where the oscillations in this function had their first local minimum. Image processing was done in ImageJ and calculation in MATLAB.

### 3. Simulation

#### a. Fundamental solution

The fundamental solution of Stokes flow (also known as the Stokeslet or Green’s function) in a liquid film was derived recently [44]. The confinement drastically changes the flow structure in these geometries, and therefore it is helpful to revise this briefly.

The hydrodynamics of an incompressible fluid at low Reynolds number are described by the Stokes equations,

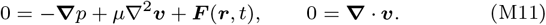

where ***v***(***r**,t*) is the flow velocity field at position ***r*** and time *t*, *p*(***r**,t*) is the pressure field, μ is the dynamic viscosity and ***F***(***r**,t*) is a force acting on the liquid. In the absence of boundaries, the flow due to a point force, ***F***_***s***_ = *δ*^3^(***r***−***r***_*s*_)***f***_*s*_, with magnitude ***f***_***s***_(*t*) and position ***r***_*s*_, with boundary conditions ***v*** = 0 as |***r***| → ∞, is

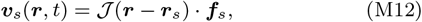

where the Oseen tensor *𝓙*_*ij*_ (***r***) in Cartesian components is

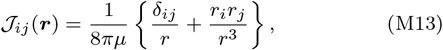

with *i, j* ∈ {*x, y, z*} and *r* = |***r***|. In a thin film, this flow is modified by the boundaries. At the bottom surface we enforce the no-slip condition, ***v*** = 0 at *z* = 0, and at the liquid-air interface we enforce the no-shear condition,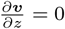 at *z* = *H*. These conditions can be satisfied exactly using a Fourier-transform method (page 56 in [44]) or approximated by truncating an infinite series of images (ibid. p. 37). The resulting expressions are rather complex, but analytically tractable. In the thin-film limit the solution simplifies to

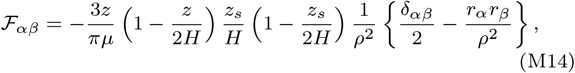

with horizontal components *α, β* ∈ {*x, y*} and relative distance *ρ*^2^ = (*x* − *x*_*s*_)^2^ + (*y* − *y*_*s*_)^2^, but vertical flows are suppressed exponentially (ibid. p. 63). Here the thin-film limit is defined as *H* ≪ *ρ*, which is equivalent to the far-field limit. The prefactors describe a half-parabolic flow profile in *z*, and the last part gives the horizontal flow structure (in curly brackets). A first difference with the Oseen tensor (Eq. M13) is that the flows decay faster, as 1/*ρ*^2^ rather than 1*/ρ*, since the surfaces impart viscous dissipation. Secondly, the streamline structure is very different.

This is demonstrated in Fig. S3, where we show the exact solution as a function of confinement, *H*, with the Stokeslet in the middle of the film, *z*_*s*_ = *H*/2. For weak confinement, we recover the bulk Stokeslet flow (Eq. M13), where the streamlines are open and oriented along the force direction. For strong confinement we recover the thin-film limit (Eq. M14), the forward flows are weaker because of surface dissipation. Moreover, a recirculation emerges with liquid moving backwards (red colours), and the streamlines all completely close onto one another. For intermediate confinement, a more complex flow structure arises with two vortexes that are separated a distance *δ* ~ *H* (green circles). Locally (*ρ* < *H*) the Stokeslet limit still holds, but globally (*ρ* > *H*) the far-field limit applies and all liquid recirculates.

Once the Green’s function 𝓕_*αβ*_ is known, it is possible to calculate the flow generated by an active carpet by integrating the Green’s function over the carpet architecture [17]. Puzzlingly however, because the liquid recirculates in the far-field in a film geometry, *the flux due to a Stokeslet through any cross-section of the film is actually zero*, for any finite film height *H*. Specifically, the flux through any *yz* plane due to a Stokeslet oriented along *x* is 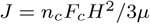. As pointed out by Liron, this is also true for channels with two parallel no-slip surfaces, or cylindrical pipes of finite radius *R*, or any double-sided confinement [27]. In an infinite or semi-infinite (single surface) geometry, however, the flux does not vanish. This apparent dilemma is resolved by realising that one may add a constant Poiseuille flow 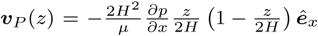 to the solution without violating the boundary conditions on the film surfaces, at *z* = 0, *H*. The flux is then specified by the pressure gradient generated by the cilia. Therefore, if each cilium on average exerts a force *F*^*c*^ along *x* and if cilia are distributed with a density *n*^*c*^ per unit area, then the flux per unit length in *y* is *J* = *n*_*c*_*F*_*c*_*H*^2^/3*μ*. Note that other forces can also generate pressure gradients, such as gravity, ***∇**p* = −*ρ****g***, which must be added to (or subtracted from) the total flux and flow profile.

#### b. Envelope approach

Here, we follow an alternative method inspired by the ‘envelope approach’ introduced by Lighthill and Blake [45, 46] to study ciliary propulsion. Instead of modelling the cilia with point forces, one can consider an envelope that covers the tips of numerous beating cilia that together form a continuous moving sheet. The no-slip condition on the bottom surface is then replaced by the motion of this envelope, ***v***_*e*_ = *U **p***, a tangential slip velocity that follows the orientation field ***p*** of the underlying cilia with an average local flow velocity *U*. This approximation is justified in the case when the cilia are close together, which is true for multiciliated cells with *N*_*cilia*_ ~ 200 cilia.

On the one hand, this approach coarse-grains the length scales smaller than the cell, so it cannot resolve the interesting flows around the individual cilia that can give rise to mixing and nutrient exchange [8, 47]. On the other hand, it is very tractable analytically (as seen in the study of micro-swimmers and active colloids [11]) and it is straightforward to implement numerically, as is discussed next, being suitable even for very large systems.

#### c. CFD solver

The flow velocity ***v***(***r***) is simulated using a 3D computational fluid dynamics (CFD) solver for the incompressible Navier-Stokes equations. This solver is optimised for low Reynolds numbers by using an implicit Crank-Nicolson method for the viscous terms and an Adams-Bashforth method for the advection terms. This algorithm is implemented on a staggered grid of size 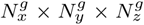 corresponding to a liquid film of size *L*_*x*_ × *L*_*y*_ × *H* [*μ*m] and with periodic boundary conditions in the *x* and *y* directions, where *x* is defined as the distal-to-proximal direction of the trachea. At the fluid-air interface we enforce the no-shear condition, 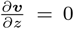 at *z* = *H*. On the tissue surface we apply the no-slip condition in the absence of cilia (*c* = 0), and in the presence of cilia (*c* = 1) we impose a slip velocity set by the envelope model, which gives the boundary condition

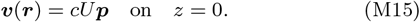

Here the cilia distribution *c*(***r***) and orientations ***p***(***r***) are either taken directly from experiments (§ 3 e), or systematically generated *in silico* for different coverage fractions, wavelengths, and disorder (§ 3 f). We simulate this system until the (unique) solution is reached at steady state, after which we save the three-dimensional velocity filed and the pressure filed of the ciliary row.

#### d. Total flux clearance time

Once the ow is solved we compute the total flux in the distal-to-proximal direction,

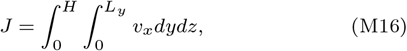

which due to incompressibility is the same for all planes perpendicular to *x*. Using the linearity of the Stokes equations, the flux can also be rewritten as

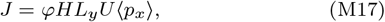

where the coverage fraction 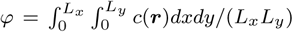. We also define the *z*-averaged flow velocity, 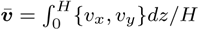, the mean longitudinal flow 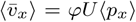, and the largest back-flow, 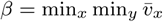.

Subsequently, the clearance time is obtained by simulating pathogen trajectories that are subject to this flow and also Gaussian fluctuations with diffusivity *D*, where the Stokes-Einstein diffusivity is *D* ~ 10^−1^ − 10^−4^ *μ*m^2^/s for micron-sized particles in mucus of viscosity *μ* ~ 10^*−*3^ − 1 Pa.s [48]. We run a Brownian dynamics (BD) simulation to find the trajectories for an ensemble of *N*_1_ ≥ 10^3^ pathogens. Initially the particle positions ***r***_*p*_ are randomly distributed across the simulation box, and follow the Langevin equation,

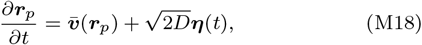

where the noise is defined by the correlation functions 〈*η*_*i*_(*t*)〉 = 0 and 〈*η*_*i*_(*t*)*η*_*j*_ (*t*^′^)〉 = *δ*(*t* − *t*^′^)*δ*_*ij*_. The Péclet number is defined as

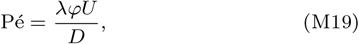

which is large if the effects of advection outcompete diffusion.

These equations are integrated with a fourth order Runge-Kutta scheme, again with periodic boundary conditions in the *x* and *y* directions. For each pathogen *i* we then record the time *T*_*i*_ taken to travel a distance *L*_*x*_ in the *x* direction, along the *D*-*P* axis. Hence, we define the clearance time *T*_*c*_ = 〈*T*〉 as the ensemble-averaged first-passage time.

Importantly, this clearance time is not only a measure of the total flux, but also of the flow structure. In particular, it is a measure of the connectivity between streamlines. If the streamlines are open and connected with the trachea outlet, the larynx, then pathogens will follow the river and rapidly be cleared. If the streamlines are closed, however, the pathogens are trapped in recirculation zones for a long time.

#### e. Experimental input

##### Large scale flows

The entire trachea is imaged with a resolution of 7176 × 4496 pixels, corresponding to 3013.9 × 1888.3 microns, with the *D*-*P* axis aligned along *x*. Here the multiciliated cells are marked with Centrin [Fig. 1B], so from this fluorescence signal we identify the positions and shapes of 29, 864 multiciliated cells. This measurement is course-grained by a factor of 5 to generate a grid of 1476 × 940 points, which subsequently is binarised to determine the coverage field *c*(***r***) = 0, 1. The orientation of the cilia cannot be measured in this assay, so for each cell we assign a stochastic direction ***p*** such that 〈*p*_*x*_〉 = 0.8, in accordance with Fig. 1H. The ciliary envelope velocity ***v***_*e*_ = *cU****p*** is then computed on each grid point of the bottom boundary in the CFD simulation. The resulting flows are shown in Fig. 2D, left panel. Note, this is not the same mouse as the one shown for the flow measurement in Fig. 2C, because that tissue is alive with beating cilia whereas the former tissue is fixed to identify the basal bodies from fluorescence.

##### Small scale flows

The tissue scale is imaged with fields of view of size 68 × 68 μ m. Here we identify the boundary of multiciliated cells by segmenting the membrane stain Vangl1 [see § 1 b], which determines the coverage field *c*(***r***). Next, we identify each cilium and its orientation by linking the relative positions of its ciliary rootlet and basal foot [see § 2 c]. For each cell, we then interpolate between its *N*_*cilia*_ ~ 200 cilia to extract the orientation field ***p***(***r***). The ciliary envelope velocity ***v***_*e*_ = *cU****p*** is then computed on each of the bottom boundary grid points of the CFD solver, with box size *L*_*x*_ = *L*_*y*_ = 68*μ*m. The resulting flow from one field of view is shown in Fig. 2D, right panel. Again, this should not be compared directly with Fig. 2C because it shows a different organism.

Nonetheless, even if it is not possible to compare the experimental and simulated flow fields directly, we compare the correlation functions and flow distributions across an ensemble of different tissues, using the same definitions as in the experiments, as described in Methods § 2 d.

#### f. Controlled variations

Next, we systematically explore how the total flux and clearance time depend on the physical properties of the mucus flow [Fig. 3A,B]. One by one, we address (1) the ciliary coverage fraction, (2) the patchiness, (3) Brownian fluctuations, (4) the spatial heterogeneity, and (5) the orientational disorder.

1. The coverage fraction *ϕ* is varied while keeping the wavelength constant, with a square periodic box of size *λ* = *L* = 128*μ*m and film height *H* = 10*μ*m. The envelope model is enforced by discretising Eq. M15 on a grid (*i, j*) of 128 × 128 cells. A patch of cilia sits at the centre of this grid, with radius 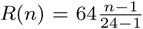 where *n* = 1, 2, 3, …, 34. The discretised coverage field is then generated with the function *c*_*ij*_ = Θ [*R*^2^ − (*i* − 65)^2^ − (*j* − 65)^2^, where Θ(*x*)is the unit step function. Consequently, the coverage fraction is *ϕ* = Σ_*i,j*_*c*_*ij*_/128^2^. All cilia are aligned along the trachea axis, *p*_*x*_ = 1, so we find the velocities 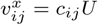 and 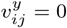at the tissue surface. The Stokes equations are solved with these boundary conditions (§ 3 c) and the resulting flows are shown in the left panels of Fig. 3C. We then simulate an ensemble of *N*_1_ = 10^4^ pathogen trajectories, initially distributed randomly, and subject to this flow and Brownian diffusion. We non-dimensionalise our system to *λ*^∗^ = 1, *U*^∗^ = 1, *D*^∗^ = 10^−4^, so the Péclet number Pé = *ϕ*10^4^, which is equivalent to a diffusivity *D* = 0.128*μ*m^2^/s, *L* = 128*μ*m and envelope velocity *U* = 10*μ*m/s. In the right panel of Fig. 3C we show the resulting mean clearance times,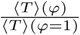, as well as the total flux, 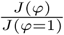, both normalised with respect to the case of full coverage (*ϕ* = 1).
2. The patchiness *λ*/*H* is explored by varying the wavelength *λ* while keeping the film height *H* = 10*μ*m constant, and also a constant periodic box size *L* = 128*μ*m, coverage fraction *ϕ* = 0.1 and cilia alignment *p*_*x*_ = 1. We again use a grid (*i, j*) of 128 × 128 cells, on which we implement a square array of *n*_*p*_ × *n*_*p*_ ciliary patches, with wavelength *λ* = *L*/*n*_*p*_ and radius 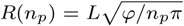, where *n*_*p*_ = 1, 2, 3, …, 24. In other words, the same number of cilia are spread out in more and smaller patches. Using these boundary conditions the flows are simulated, which are shown in the left panels of Fig. 3D. For large patchiness a back flow emerges. We again simulate *N*_1_ = 10^4^ pathogen trajectories with constant diffusivity *D* = 0.128*μ*m^2^/s, *L* = 128*μ*m and mean flow velocity 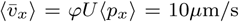. In the right panel of Fig. 3D we show the resulting mean clearance times, 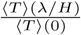, as well as the to-tal flux, 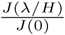, and the back flow, 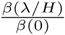, all normalised with respect to the case of uniform coverage (*λ* = 0).
3. The geometric order is varied by considering the crystallinity *γ*, while keeping the coverage fraction *ϕ* = 0.1, the mean patchiness 〈(*λ*)〉/*H* = 8 and the cilia alignment *p*_*x*_ = 1 constant. We begin with a crystalline array of 4 *×* 4 cilia patches, as in § 3 f*2* above with *n*_*p*_ = 4. These patches are displaced spatially by a random vector (*X, Y*), where both components are drawn from a Gaussian distribution with standard deviation *σ*. The geometric order parameter is then defined as 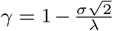, the crystallinity. As before we implement these patches on a periodic 128 × 128 grid, but we also consider periodic image systems so that if (*X, Y*) leaves the grid, it appears on the other side. Moreover, we allow patches to overlap randomly, so the relation 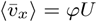 remains satisfied. We generate these initial conditions for an ensemble of *N*_2_ = 50 epithelia, and for *N*_3_ = 16 different values of the order parameter, so *N*_2_*N*_3_ = 800 flows are simulated. A few examples of these flow patterns are shown in Fig. 4A. For each configuration we then simulate *N*_1_ = 1000 pathogen trajectories, with a Péclet number of Pé = 10^3^, so 800,000 trajectories total. In the right panel we show the resulting clearance times, now averaged over the ensembles *N*_1_ and *N*_2_, 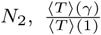, and the total flux, 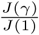, both normalised with respect to the case of crystalline order.
4. The effect of cilia alignment is examined by varying the orientational order parameter, *M* = 〈*p*_*x*_〉, while keeping the coverage fraction *ϕ* = 0.1, the patchiness *λ*/*H* = 8 and the crystallinity *γ* = 1 constant. Again we begin with a crystalline array of 4 *×* 4 cilia patches, *c*_*ij*_, as in § 3 f*2* with *n*_*p*_ = 4. Now each cilia patch *k* is given a random orientation ***P***_*k*_, which is drawn from a circular Gaussian, the Von Mises distribution,

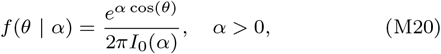

where *α* is the distribution width and the Bessel function of the first kind *I*_0_(*α*) is just a normalisation factor such that ∫*f* (*θ*)*dθ* = 1. Random numbers *θ* are drawn from this distribution using the Smirnov transform, i.e. solving the inverse of the cumulative distribution, such that 〈cos(*θ*) = *M* ϵ [0, 1]. Hence we compile a grid for the longitudinal and perpendicular surface velocities, 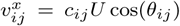 and 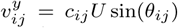, where *θ*_*i,j*_ = *θ*_*k*_ for each patch. These initial conditions are generated for an ensemble *N*_2_ = 50 epithelia, and for *N*_3_ = 16 different values of the order parameter, 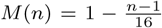. We again simulate *N*_1_ = 1000 pathogen trajectories for each configuration, with Pé = 10^3^. In the right panel of Fig. 4B we show the resulting clearance times averaged over *N*_1_ and *N*_2_, 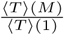, and the total flux, 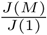, both normalised with respect to the case of aligned cilia.
5. The effect of Brownian fluctuations is tested by varying the Péclet number, while keeping constant the wavelength *λ* = *L* = 128*μ*m, the film height *H* = 10*μ*m, the coverage fraction *ϕ* = 0.1, crystallinity *γ* = 1, and cilia alignment *p*_*x*_ = 1. We use the same ciliary arrangement 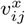 as before, (§ 3 f*2* above with *n*_*p*_ = 1), for which the flow is shown in the fourth panel of Fig. 3D. Using this flow, in Fig. 4C the trajectories of 50 particles are presented for each value of the Péclet number, Pé = 100, 100, 10, by varying the diffusivity *D*. We then gather statistics of *N*_1_ = 10^4^ trajectories for the Péclet number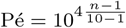, where *n* = 1, 2, 3, …, 10. In the right panel we show the resulting mean clearance times, 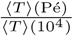, and the total flux, 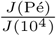, both normalised with respect to the case of weak fluctuations.

#### g. Phase diagrams

Finally, we cross-compare these parameters in three phase diagrams that combine the coverage fraction, the patchiness, and the ciliary orientation. For all diagrams, we generate an ensemble of ciliary carpet structures starting from a crystalline array of *n*_*p*_ × *n*_*p*_ cilia patches, as in § 3 f*2* with *γ* = 1. As before the coverage fraction is varied by changing the patch size *R* so that 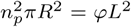, the patchiness is varied by the number of patches *n*_*p*_ and the patches are given a ciliary orientation with order parameter *M* = 〈*p*_*x*_〉 using the Von Mises distribution.

1. For the *ϕ − λ* diagram we simulate *N*_2_*N*_3_ epithelia for *N*_2_ = 14 values of coverage and *N*_3_ = 10 values of patchiness. In the resulting flow profiles we simulate *N*_1_ = 10^3^ tracer trajectories with Pé/*ϕ* = 10^4^ and constant *p*_*x*_ = 1, which together yield the mean clearance time. The clearance time is normalised with respect to an ideal carpet, *ϕ* = 1 and *λ* = 0.
2. For the *ϕ*−*M* diagram we simulate *N*_2_*N*_3_*N*_4_ flows for *N*_2_ = 14 different values of coverage, *N*_3_ = 16 different values of orientational order, and for each an ensemble of *N*_4_ = 20 epithelia. In all the resulting flow profiles we simulate *N*_1_ = 10^2^ tracer trajectories with Pé/*ϕ* = 10^3^ and constant *λ*/*H* = 8. The clearance time is normalised with respect to *ϕ* = 1 and *M* = 1.
3. For the *λ*−*M* diagram we simulate *N*_2_*N*_3_*N*_4_ flows for *N*_2_ = 16 different values of patchiness, *N*_3_ = 14 different values of orientational order, and for each an ensemble of *N*_4_ = 20 epithelia. In all the resulting flow profiles we simulate *N*_1_ = 10^2^ tracer trajectories with Pé = 10^4^ and constant *ϕ* = 0.4. The clearance time is normalised with respect to *λ* = 0 and *M* = 1.

### 4. Video captions

#### Video 1. Activity of a field of multiciliated cells in the trachea

Time-lapse ex-vivo DIC imaging of a section of the mouse trachea multiciliated tissue.

#### Video 2. Ciliary driven flows are globally coherent

Flow generated by multiciliated cells across the entire trachea. Flow is imaged by time-lapse confocal microscopy. Fluorescent beads serve as tracer particles.

#### Video 3. Ciliary driven flows are locally heterogeneous

Microstructure of the flow generated by a subsection of the trachea multiciliated tissue. Flow is imaged by time-lapse confocal microscopy. Fluorescent beads serve as tracer particles.

#### Video 4. Activity of multiciliated cells in the trachea

Time-lapse ex-vivo DIC imaging of a cross-section of the mouse trachea multiciliated tissue.

#### Video 5. Fluctuations improve particle clearance in patchy arrangements of multiciliated cells

Particle trajectories (*N*_*p*_ = 1000) are simulated for a range of noise strengths. Initially, the particles are distributed evenly and identically in all three panels, emulating a uniform deposition from breathed-in air. Over time, they are subject to flow (same as in Fig. 4C) and fluctuations with different diffusion constants, such that the Péclet number is Pé = 10^4^, 10^3^, 10^2^, respectively. Colours indicate particle index and black trails show their pathlines. For weak noise the particles accumulate in recirculation zones, but strong fluctuations facilitate a faster clearance.

## Supporting information

Video 1. Activity of a field of multiciliated cells in the trachea

Video 2. Ciliary driven flows are globally coherent

Video 3. Ciliary driven flows are locally heterogeneous

Video 4. Activity of multiciliated cells in the trachea

Video 5. Fluctuations improve particle clearance in patchy arrangements of multiciliated cells

## ACKNOWLEDGEMENTS

This work was supported by funding from the National Science Foundation Center for Cellular Construction (NSF grant DBI-1548297), a Human Frontier Science Program Fellowship to A.M. (LT001670/2017), and a Ruth L. Kirschstein National Research Service Award to M.H. (F32HD089639). We thank Kathryn Anderson (Sloan Kettering Institute) for providing the Arl13bmCherry/Centrin-GFP animals used for this study and the Stanford Research Computing Center for providing computational resources and support. We thank William Gilpin for providing comments on the manuscript.

## AUTHOR CONTRIBUTIONS

GRS, WM and MP designed the research. GRS and MH performed the tissue imaging. GRS analysed the data. AM contributed intellectually to the paper and developed the simulations. GRS and AM wrote the manuscript. MH and LJ provided key reagents and resources for tissue imaging. All authors edited the final manuscript.

## COMPETING INTERESTS

The authors declare no competing financial interests.

## DATA AVAILABILITY

The data that support the plots within this paper and other findings of this study are available from the corresponding author upon request.

## CODE AVAILABILITY

The computer codes used in this paper are available from the corresponding author upon request.

---

**FIG. S1.**
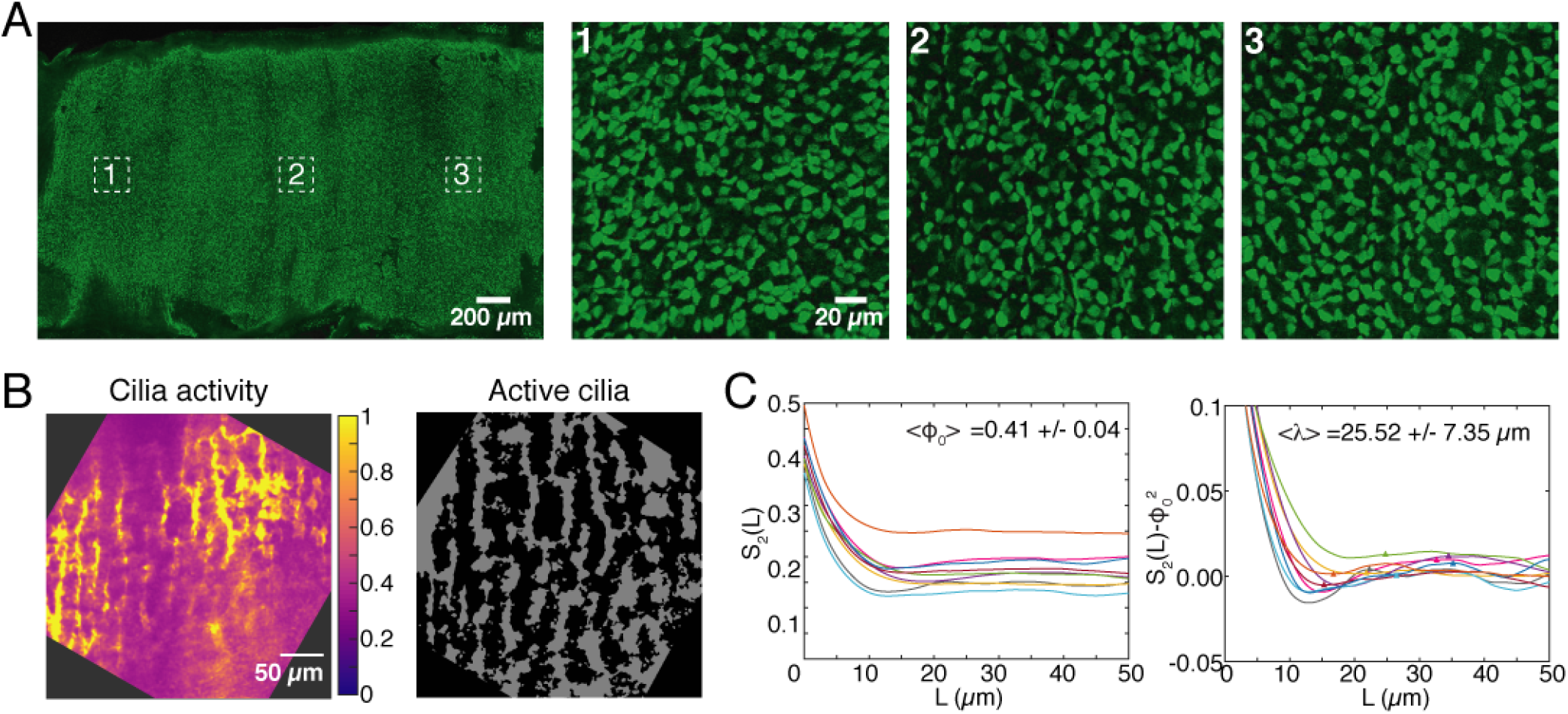
**A.** Immunofluorescence image of a trachea expressing Centrin-GFP (left). Regions outlined by the squares are magnified to the right, showing that multiciliated cells form a “patchwork” pattern in the distal (1), medial (2) and proximal (3) regions of the trachea. **B.** Heat map scaled such the 1 corresponds to the regions of maximum cilia activity in Supp. Video 1. Right: Binary image where white regions show regions where cilia are active in Supp. Video 1 (See Methods § 2 b). **C.** Average coverage fraction and wavelength measured by calculating the two point correlation function of from binary images where regions with cilia activity were identified (e.g. image shown in B).

**FIG. S2.**
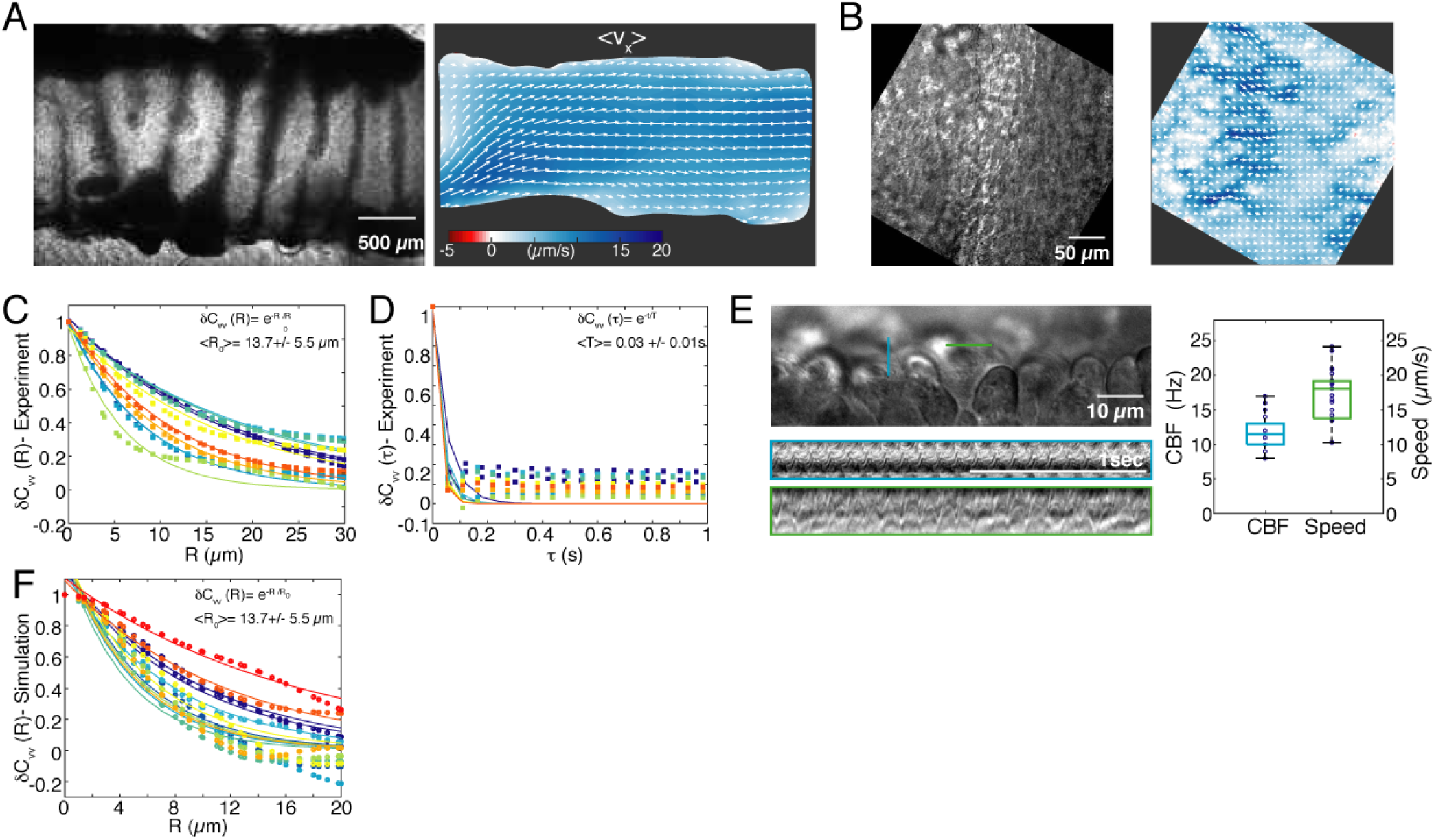
**A.** Transmitted light image of trachea showing the scale at which flow is measured. Right: Longitudinal flow strength (*v*_*x*_, red-blue) generated by trachea pictured left. Entire flow strength and streamlines shown in (Fig. 2A,C; left). **B.** Representative transmitted light image of a typical region of the tissue where flow microstructure is analysed. Right: Longitudinal flow strength (*v*_*x*_, red-blue) generated by trachea pictured left. Entire flow strength and streamlines shown in (Fig. 2A,C; right). **C.** Plot of spatial correlations of the flow field. Each trace is the correlation function M9 calculated for a field of view of the size shown in B. Points correspond to values measured, solid line shows exponential fits. **D.** Plot of temporal correlations of the flow field. Each trace is the correlation function M10 calculated for a field of view of the size shown in B. Points correspond to values measured, solid line shows exponential fits. **E.** Quantification of beat frequency and tip velocity of cilia in multiciliated cells. Kymographs drawn from the lines shown in DIC image. Each peak corresponds to one cilia beat cycle (blue), while the slopes of the line indicate the rate at which cilia tips move over time (green). **F.** Plot of spatial correlations of simulated flow fields. Measurements of multiciliated cell organisation were used as an input for these simulations (See Methods § 3 e). Each trace is the correlation function M9 calculated for a simulated flow field. Points correspond to values calculated, solid line shows exponential fits.

**FIG. S3.**
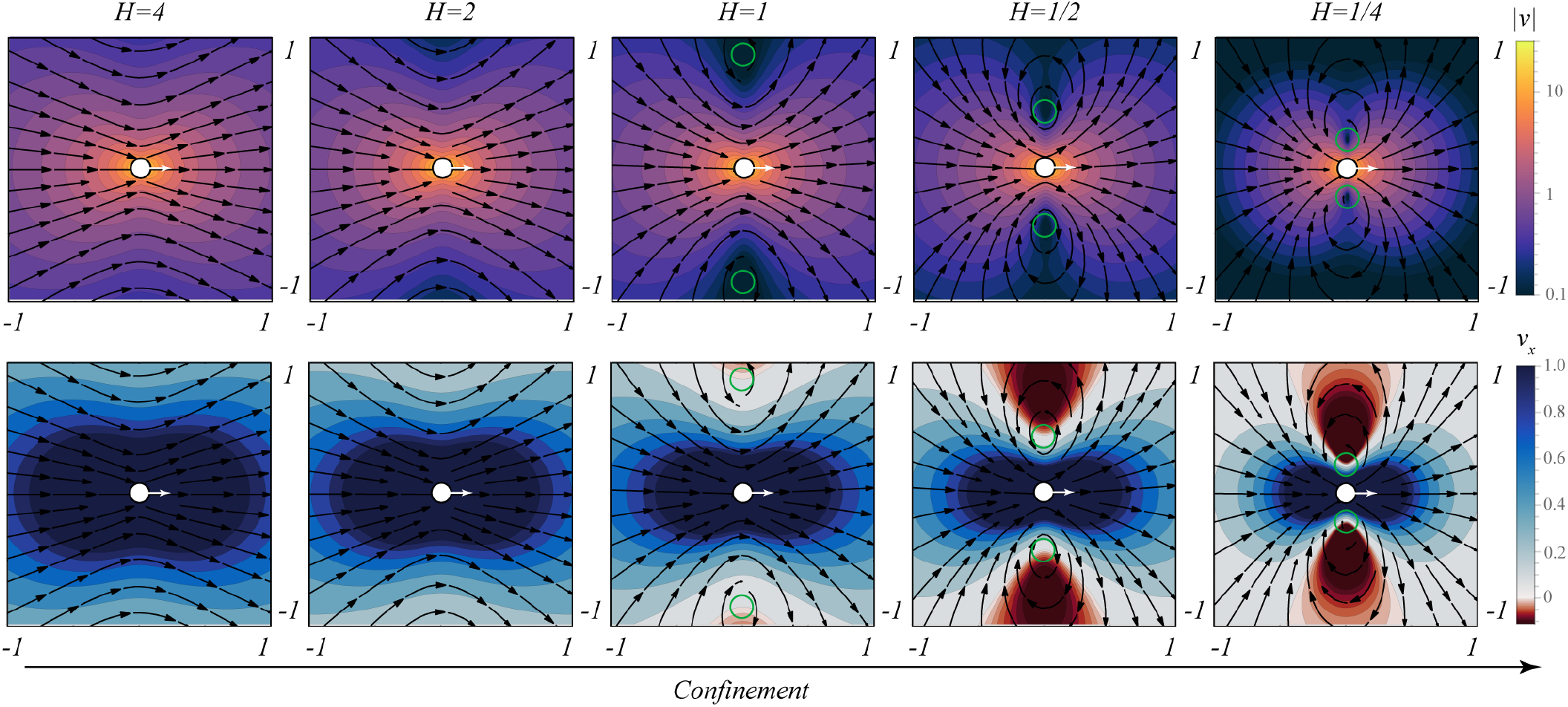
Effect of confinement on Stokeslet flow in a liquid film. The film height in the *z* direction is *H*, compared to the horizontal scale *x, y* ∈ [−1, 1]. The Stokeslets are located in the middle of the film, at *z* = *H*/2, and oriented in the *x* direction (white arrows). We use 9 image reflections, (*n*) = (0) − (9) in Table 1 of [44]. **Top** panels: Magnitude (thermal) and streamlines (black arrows) of the flow velocity. **Bottom** panels: Same, showing the *x* component (red-blue) of the flow. For thin films a recirculation emerges, with a vortex centre marked in green.

**FIG. S4.**
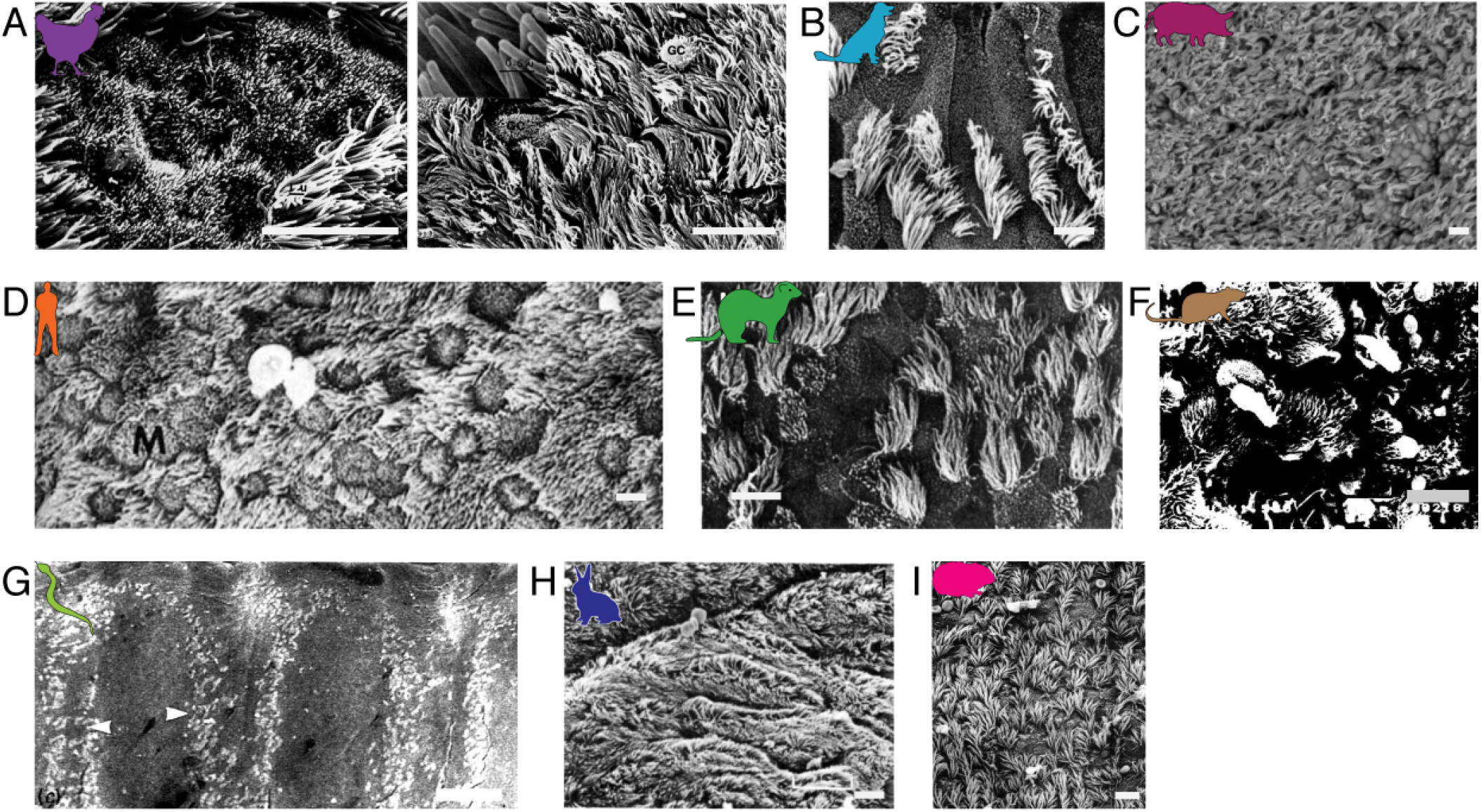
From the literature: SEM images of airway multiciliated tissue. **A.** Chicken; Image adapted from [36]. **B.** Dog; Image adapted from [37]. **C.** Pig; Image adapted from [41]. **D.** Human; Image adapted from [40]. **E.** Ferret; Image adapted from [38]. **F.** Rat; Image adapted from [42]. **G.** Snake; Scale bar=100*μm*. Image adapted from [43]. **H.** Rabbit; Image adapted from [49]. **I.** Hamster; Image adapted from [39]. Scale bar=10*μm* for all panels except G.

